# Flow parsing as causal source separation: A computational model for concurrent retrieval of object and self-motion information from optic flow

**DOI:** 10.1101/2024.11.04.621864

**Authors:** Malte Scherff, Markus Lappe

## Abstract

Optic flow, the retinal pattern of motion that is experience during self-motion contains information about one’s direction of heading. If the visual scene contains other moving objects optic flow becomes a combination of self-motion and independent object motion. The global pattern due to self-motion is locally confounded, and the locally restricted object flow is the sum of components due to these different causal sources of motion. Nonetheless, humans are able to retrieve information from such flow accurately, including the direction of heading and the scene-relative motion of an object.

One way to handle such complex flow is flow parsing, a process speculated to allow the brain’s sensitivity to optic flow to separate the causal sources of retinal motion in information due to self-motion and information due to object motion. In a computational model that retrieves object and self-motion information from optic flow, we implemented such a process of causal separation based on heading likelihood maps, whose distributions indicate the consistency of parts of the flow with self-motion alone. The flow parsing allows for concurrent estimation of heading, the detection and localization of and independently moving object, and the estimation of scene-relative motion of that object.

We developed a paradigm that allows the model to perform all the different estimations while systematically varying how the object contributes to the flow field. Simulations of that paradigm showed that the model replicates many aspects of human performance, including the dependence of heading estimation on object speed and how different object movements bias that estimation. Regarding object detection and motion estimation, the model’s results fit human behavioral data, the latter even for flow of reduced quality.

## 1. Introduction

Optic flow is the pattern of light on the retina that results from relative motion between an observer and their visible surroundings. It provides valuable information about the structural layout of the scene, including its rigidity, and the different types of motion present (Gibson, 1950; Longuet-Higgins and Prazdny, 1980). Recovering self-motion information from optic flow is not only theoretically possible but also crucial for safe navigation in daily life (Warren et al., 1988; Cutting et al., 1992; Lappe et al., 1999). Likewise, detecting an independently moving object (Royden and Connors, 2010; Royden and Moore, 2012), estimating its trajectory (Rushton and Warren, 2005; Warren and Rushton, 2007, 2008, 2009) and determining its time to contact (Gray et al., 2004; Gray and Regan, 2000) during self-motion is evident when considering simple examples in sports. For instance, a soccer player must steer their movement towards a position on the pitch where the ball will likely land to receive an inaccurate pass from a teammate successfully.

However, it remains unclear how the two processes - estimating self-movement and deriving information about sources of independent motion - are connected and interact when they are based on the same flow information, and how the causal attribution of any motion to self-motion or object motion is achieved.

### 1.1. Theoretical considerations and psychophysical evidence for the estimation of self-movement

The simplest form of flow patterns is one in which everything moves radially away from a singular point, the focus of expansion (FOE). Such a radial flow field occurs when an observer moves through a stationary scene, and the movement solely consists of a translation in a certain direction. In that case, the direction of self-motion and the FOE coincide. It is well established that humans can recover the direction of motion from such flow patterns with an accuracy well within a range of 1 to 2 degrees of error (Warren and Hannon, 1988; Warren et al., 1988; Cutting et al., 1992).

While such radial patterns are often used in psychophysical studies, it is well known that eye-movements that occur during self-motion confound this simple structure by adding rotational components (Regan and Beverley, 1982a; Warren and Hannon, 1990; Lappe et al., 1998; Calow and Lappe, 2007; Matthis et al., 2022). Nonetheless, studies with real as well as simulated eye movements showed that self-motion estimation is still possible (Warren and Hannon, 1990; Royden et al., 1992, 1994; Li and Warren, 2000). Further studies have shown that self-motion estimation is quite robust such that adding noise to flow stimuli reduces but does not impede the ability to estimate the direction of self-motion (Warren et al., 1991; Van Den Berg, 1992).

Studies have also shown that the presence of an independently moving object (IMO) in the optic flow can bias heading estimation (Warren and Saunders, 1995; Royden and Hildreth, 1996; Layton and Fajen, 2016b; Li et al., 2018). In some of those studies, the objects in the scene moved such that the combined flow only consisted of lateral motion. By moving the object backward the same amount the observer moved forward the relative motion between IMO and the observer was only due to the object movement. Hence, the combined flow of the IMO contained no information about the heading direction. Heading estimation was then biased in the direction of the object movement (Royden and Hildreth, 1996; Li et al., 2018). Otherwise, when the combined flow of the IMO was the result of object and observer movement, the bias was in the opposite direction of the object motion (Warren and Saunders, 1995; Layton and Fajen, 2016b,d; Li et al., 2018). Another observation about the influence of an IMO on heading estimation was made in a study in which the object’s speed was systematically varied (Dokka et al., 2019). Increasing the speed of the object from zero first led to an increase in heading estimation error, which peaked for the intermediate speeds tested, and then reduced the object’s impact on the heading estimation as the error dropped to nearly zero for further increases of object speed.

### 1.2. Theoretical considerations and psychophysical evidence for the estimation of independent object motion during self-motion

Gaining information about an IMO while moving from optic flow alone presents a challenge, as no part of the visual field exclusively contains image motion caused by the object’s movement. Selfmotion always confounds it. Nonetheless, humans are able to detect an IMO in an optic flow stimulus based on the divergence in direction (Royden and Connors, 2010) or speed (Rushton et al., 2007; Royden and Moore, 2012) to the rest of the pattern. When participants were tasked with judging the trajectory of an IMO in a radial flow field, the results were consistent with an interpretation in a world-relative reference frame (Rushton and Warren, 2005; Warren and Rushton, 2007, 2008, 2009). These results align with the idea of a process called *flow parsing* with which the brain separates retinal motion components due to self-motion from those due to object motion. While no explicit mechanism was proposed, this might be achieved by using the brain’s selectivity to flow patterns (e.g. Duffy and Wurtz (1991)) to identify and globally discount the component due to the self-movement. The remaining flow could then be attributed to an indecent source of motion and used to gain information about it. This might amount to an iterative solution, in which self-movement estimation (possibly biased) is followed by determining object movement, which relies on the prior estimate of self-movement, and then the process may be iterated to refine each solution (Pauwels et al., 2010; Layton and Fajen, 2016b).

However, some studies have, seemingly paradoxically, suggested that estimation of independent object motion does not rely on prior estimation of selfmotion. For example, Warren et al. (2012) showed that participants’ judgment of an object’s trajectory was not consistent with its scene-relative movement when self-motion perception was strategically biased (Warren et al., 2012) and proposed that flow parsing might occur independently of heading estimation. Further evidence for that was provided by (Rushton et al., 2018a) as they found performance in object movement estimation to be more precise than in heading estimation. Therefore, it is unlikely that the identification of scene-relative object movement relies on the prior self-motion estimation, especially as another study suggests that such identification might solely be driven by optic flow processing (Rushton et al., 2018b). These findings suggest that self-motion and object motion estimations might share initial processing stages but might ultimately take place in parallel and independent of each other.

### 1.3. Aim of the study

We present a computational model that processes optic flow and can estimate self-motion direction and parameters of an IMO in parallel. To do so, the model uses flow parsing to separate information from different causal sources of flow. The flow parsing process is based on likelihood maps computed for different parts of the flow field. These maps indicate the consistency of the corresponding flow with various heading directions. Depending on that consistency, the likelihood maps are used for either heading or object estimation. The model’s structure allows these estimations to be run in parallel without needing recurrent or feedback connections.

When comparing the model’s performance for a self-motion scene, including an IMO, with research from studies in the literature, the results align with behavioral data. To be more specific, the heading estimation process gives rise to an error that systematically depends on object speed, with slow and fast moving objects causing a small error and an error peak for intermediate speeds, similar to findings of Dokka et al. (2019). Additionally, the direction of mis-estimation depends on whether the object maintains a fixed depth relative to the observer, a finding reported in various studies (Warren and Saunders, 1995; Royden and Hildreth, 1996; Li et al., 2018; Layton and Fajen, 2016b,d). The object’s detectability depends on the deviation of the object flow from the background pattern, as Royden and colleagues found (Royden and Connors, 2010; Royden and Moore, 2012). While there is no research providing behavioral data in regard to object localization in optic flow fields, the model is able to successfully localize the independent source of motion solely based on flow velocities. Lastly, the object direction estimation is similar to human performance that shows that the perceived trajectory is consistent with scene-relative motion (Warren and Rushton, 2007, 2008, 2009).

### 2. The computational model

Mathematically, optic flow arising from self-motion in a static scene is described by a set of flow vectors. These vectors represent the image velocity, that is, the time derivative of the spatial components of the projection of the 3D points in the environment onto the image plane. This plane is placed at distance *f*, perpendicular to the line of sight, and acts as a simplified representation of the retina of the monocular observer. The relationship between the instantaneous observer movement and the optic flow field can be described with the following equation:

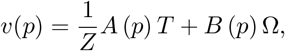

that describes the image velocity for a static scene point *P* = (*X, Y, Z*), with *p* = (*x, y*) = *f* (*X/Z, Y/Z*) as the image coordinates, and *T* and Ω as the motion parameter of the observer, the translation direction and rotational velocity, respectively (Longuet-Higgins and Prazdny, 1980; Heeger and Jepson, 1992).

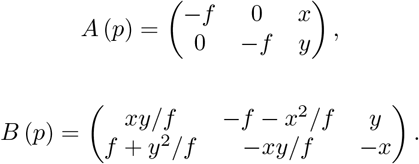

Written in this form, this equation illustrates that optic flow fields consist of two components, where only the translational part, not the rotational part, depends on the depth. If P belongs to a non-static scene object with *S* describing its translation direction in the world, the equation can still be used to compute the respective flow. For Ω = (0, 0, 0) it holds:

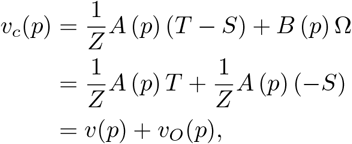

describing the combined flow due to the relative motion (T-S) between the observer and the object, with *v*_*O*_ as the object flow.

### 2.1. Subspace Algorithm

Trying to find the best self-motion parameter to explain a given flow field imposes the challenge of solving a set of equations with many unknowns: the 6 self-motion components and depth values of every point in the scene that contribute to the flow field. While there are different approaches to solving this problem, this work focuses on a particular method, the subspace algorithm developed by Heeger and Jepson (1992). This method allows for successive solving for the observer translation direction, then the rotation, and finally the depth structure of the scene based on the prior results. The relevant part for this study is the first one. To solve for the translation, a type of residual function is defined that calculates how consistent a candidate translation direction is with the given flow vectors, allowing for a least-square estimation approach by picking the candidate direction that minimizes the residual value. To define said residual function for a candidate direction *T* and given flow *v* at image locations *p*_1_ = (*x*_1_, *y*_1_), …, *p*_*n*_ = (*x*_*n*_, *y*_*n*_) we define

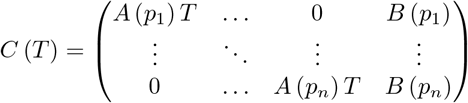

with the matrices as defined above. The residual function is now defined as

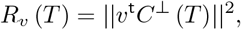

with *C*^⊥^ (*T*) depicting the orthogonal complement to *C* (*T*). Sampling the set of all candidate translation directions, the heading space, in a retinotopically organized way and evaluating the corresponding residual functions on it yields a surface of residual values that maintains the same retinotopic organization. Such a surface can be seen as a heading likelihood map.

### 2.2. Implementations

Since its inception, the subspace algorithm has been implemented in different heading estimation models. Heeger and Jepson (1992) as well as Sauer et al. (2022) used straightforward implementations that evaluated the residual functions on flow fields based on different self-motion scenarios. The former tested their model on artificial flow fields derived from simulated 3D scenes and realistic ones from real-world camera recordings (Heeger and Jepson, 1992). Compared to other heading estimation methods available then, their model performed well and recovered translation even when varying degrees of uniform noise were present in the flow. Sauer et al. (2022) used the algorithm to estimate heading for distorted optic flow in simulated selfmotion scenarios that incorporated the effects of ophthalmic correction lenses. The strength of the distortion effect on the flow varied depending on where the gaze crossed the lens. These distortions acted as a continuous, systematic noise across the visual field, altered the perceived flow, and resulted in a heading estimation bias.

Lappe and Rauschecker (1993, 1995b) devised a biologically plausible implementation of the subspace algorithm in a two-layer neural network that estimates heading from optic flow fields. The input layer, designed after monkey area MT, contains neurons that encode motion direction and speed, which results in a population-encoded representation of the optic flow. Neurons in the second layer are selective for directions of translational egomotion and model the next stage of the flow processing pathway, area MSTd. The activities of populations of these neurons form a retinotopic heading map, which corresponds to the residual function in which the activity peak indicates the direction most consistent with the input flow. To achieve this output, the connection strengths between the neurons of the two layers are precomputed using the subspace algorithm. Heading estimation performance was tested across a variety of settings with this model (Lappe and Rauschecker, 1993, 1994; Lappe, 1996, 1998; Lappe et al., 1999). Results were well in line with human behavior as long as the input flow was dense enough, i.e., it consisted of at least 10 points. When the 3D scene was sufficiently non-planar, or the level of uniform noise added was low enough, even simulated eye-movements did not significantly interfere with the heading estimation. Additionally, the model could reproduce the mislocalization of the FOE under the optic flow illusion (Lappe and Rauschecker, 1995a).

### 2.3. Structure of residual surfaces

All those implementations have in common that only the peaks of the computed surfaces were used for the final heading estimations, and no further attention was paid to the distributions of the residual values. Hence, only some of those distributions were reported for illustration purposes. While Heeger and Jepson showed a residual surface with two peaks, indicating multiple distinct translation directions as a potential solution, they clarified that the corresponding flow field was ambiguous. Various translations towards distinct planes can result in similar flow fields (Longuet-Higgins and Prazdny, 1980; Regan and Beverley, 1982b; Grigo and Lappe, 1999). Apart from that, only surfaces with a single peak were reported by the corresponding authors, and we found them when examining similar flow fields and the resulting residual distributions as well. All flow fields tested were due to self-motion through a rigid environment. On the other hand, multiple peaks are often found when the scene includes an independent source of motion.

Fig.1 shows residual distributions for three flow fields. The first distribution is based on flow solely due to observer translation. Hence, the surface has a single peak. The structure of the residual distributions changes when object motion is introduced into the scene, as seen in the second panel. There, the flow field contains an object placed to the right of the FOE. The object moves such that the combined flow is lateral to the right and mostly in line with the overall flow pattern. In terms of speed, the combined flow is faster than the observer flow. This results in a residual surface in which a second peak emerges compared to the first surface. The area between the peaks resembles a saddle point as it is also a local minimum in the direction orthogonal to the peaks, and the saddle point’s position coincides with the IMO’s location. The last panel shows an example of a residual surface for when the combined flow is faster than the surrounding observer flow and deviates in direction from the flow pattern. The combination of observer and object movement gives rise to purely vertical combined flow, leading to a residual surface with a distinct saddle point, again matching the object’s location. This time, the saddle point’s orientation, that is, the axis on which the peaks emerge, does not entirely align with the combined flow direction but is slightly tilted relative to vertical.

**Figure 1:**
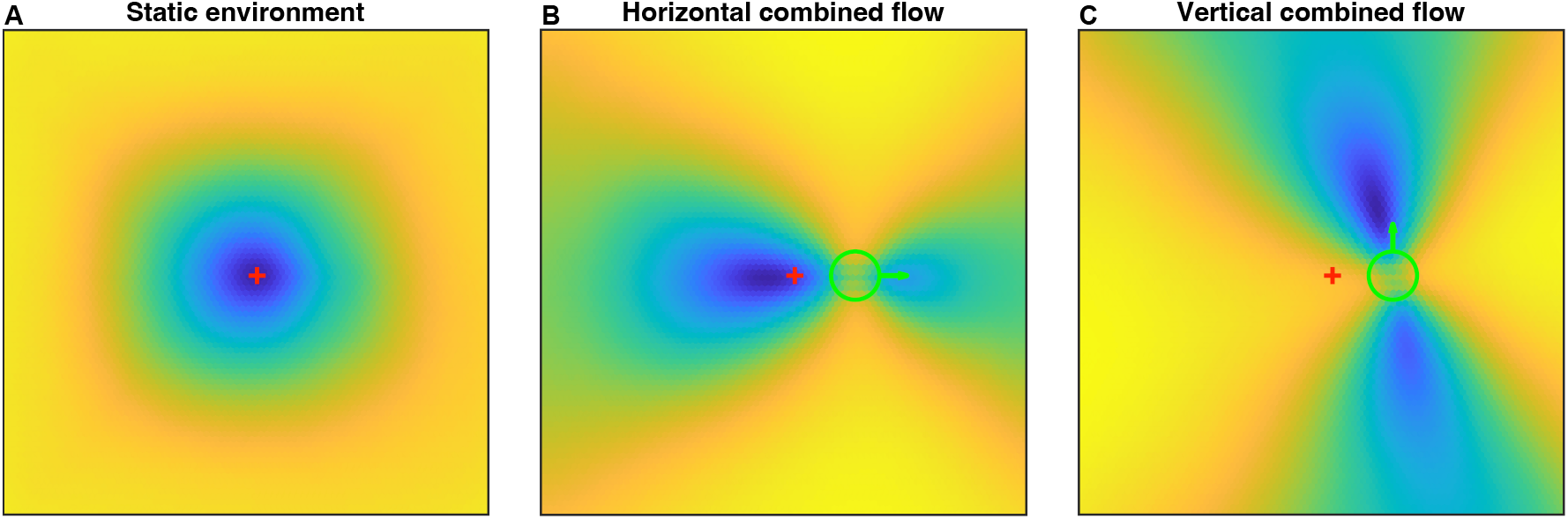
Examples of residual surfaces. Corresponding flow fields are due to simulated observer translation towards a static cloud of dots. The red cross indicates the FOE of the radial flow pattern and, therefore, the heading direction. The object, if present in the scene, is depicted with a green circle, and the green arrow indicates the direction of the combined flow. The residual surface has a singular peak, which coincides with the true heading direction when no object is present (A). When present, a moving object gives rise to residual surfaces with a saddle point at the object’s location, and the position of the surrounding peaks changes with the object’s flow direction (B-C).

Overall, the residual structures carry information about the independent source of motion in the scene. The retinal location of the object is consistently indicated by the saddle point between the residual peaks, and its orientation of the peak placements varies depending on the combined flow direction.

### 2.4. Model

The computational model we present here aims to perform three tasks: infer the causal sources of retinal motion, estimate self-motion and estimate the parameters of an IMO. The first task is implemented via a flow parsing process, which is based on the aforementioned difference in residual distributions for flow fields with or without an IMO. This process then regulates the inputs to the self-motion and IMO processes to minimize the use of information from flow not relevant to the respective tasks. The multi-layered model only contains forward connections between the layers, so neither process is influenced by any outcome.

In most of the layers, we employ a specific type of operator to process the input the layer receives. All types of operator are at least loosely inspired by certain types of neurons and their capabilities as well as earlier implementations of the subspace algorithm. As input from and output provided by each layer are represented as retinotopic maps and the range of each operator is restricted to a certain area of these maps, we refer to these areas as the receptive fields of the operators. The general structure of the model, as well as a more detailed presentation of all layers, can be seen in Fig.2.

**Figure 2:**
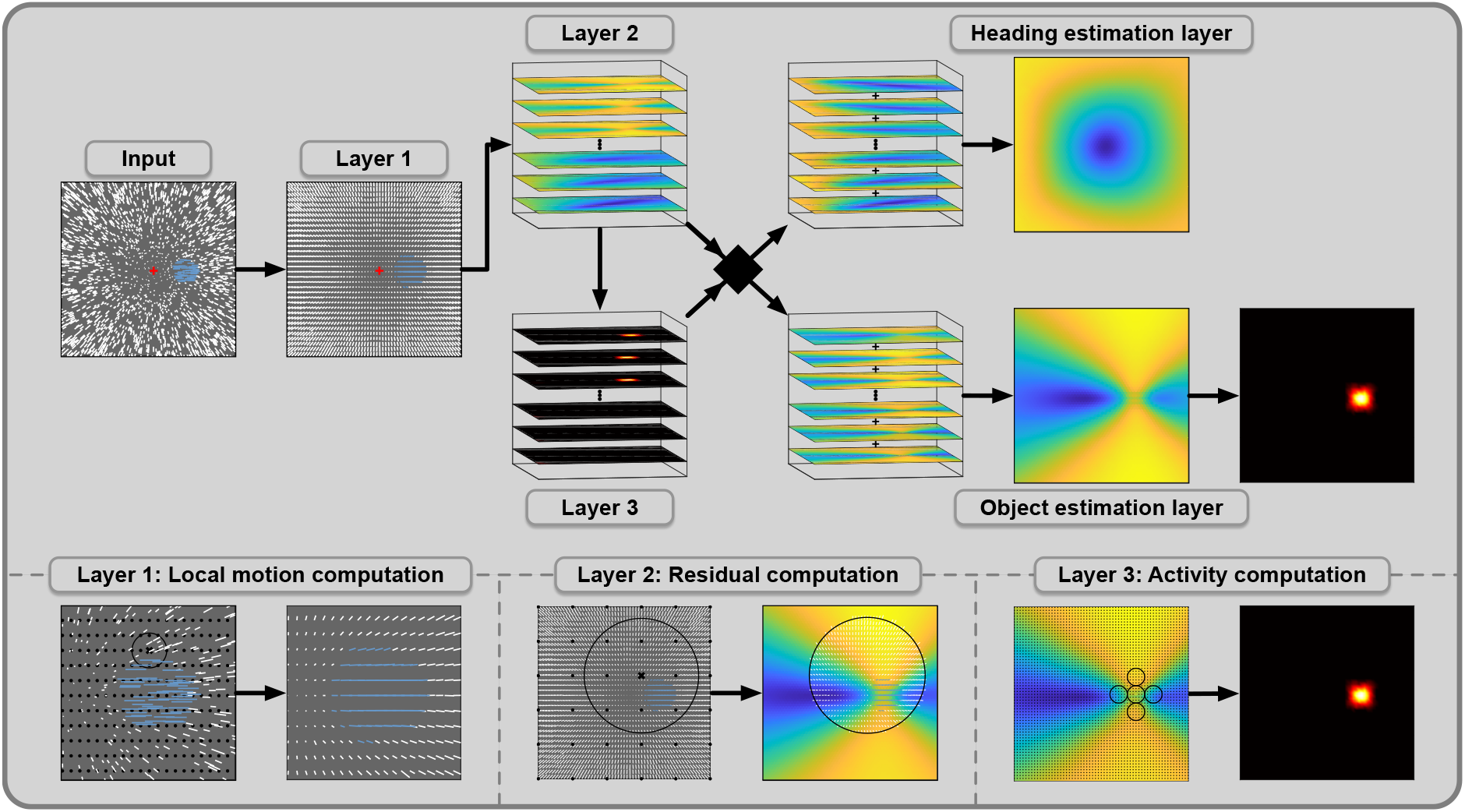
Structure of the computational model. The model processes optic flow to detect the presence of a moving object, estimate self-motion direction, estimate object motion direction, and determine object location. It consists of multiple layers with specific processing operators in each layer. Optic flow fields form the input into the model. White and blue vectors represent observer flow and combined flow, respectively. Layer 1 computes a vector representation of the input flow by computing local averages of speed and direction. It contains operator units with small circular receptive fields that cover the FOV. The black circle shows the receptive field of one operator as an example, and the locations of the remaining operators are indicated with black dots. Layer 2 computes residual surfaces for flow in different parts of the visual field. Flow vectors in the receptive field of one operator are grouped, and one residual surface is computed for each group. Layer 3 computes activity maps, one for each residual surface. Operators calculate average residual values in parts of their cross-shaped receptive fields and use differences between them to compute activity. Receptive fields vary in size and orientation and cover the residual surface. The flow parsing process is based on the activity maps and decides whether a residual surface is used for heading or object estimation. The corresponding surface is used for object estimation when the activity maximum is high enough. The heading estimate is determined by the peak of the heading map, which is the result of summing the respective surfaces. For the object estimation, the incoming surfaces are summed up, and an activity map is computed. Size and position of the activity maximum determine object detection and localization, respectively. To estimate the object’s movement direction, the orientation of the operators that contributed to the activity is used.

The goal of the first layer is to create a vector representation of the incoming optic flow by averaging the flow at different retinal locations (Lappe, 1996; Calow et al., 2005). Based on neurons found in the middle temporal visual area (MT) that show selectivity for direction and speed (Albright, 1984; Maunsell and Van Essen, 1983), we place the operators of this layer so that the field of view (FOV) is covered by the corresponding receptive fields in which the mean speed and direction of flow are computed. This results in a vector representation of the optic flow across the FOV.

Area MT projects to the dorsomedial region of the medial superior temporal area (MSTd) that contains neurons showing heading-sensitivity based on larger flow patterns typical for self-motion (Duffy and Wurtz, 1991). Hence, the second layer aims to test the consistency of flow from different retinal regions with the candidates in the heading space Lappe and Rauschecker (1993); Lappe et al. (1996). The retinal regions are defined by the receptive fields of the operators corresponding to this layer that cover the visual field. Flow vectors in the same receptive field are grouped, and residual functions defined by the subspace algorithm are evaluated on each group. This results in residual surfaces, one for each of the groups.

The third layer plays a crucial role in the model, serving as the foundation for the flow parsing process. It assesses the distribution of the residual values since a distinct saddle point indicates the presence of a source of independent motion in the corresponding retinal region. Containing one, such surfaces will be used for the object estimation process, while single peak surfaces will contribute to the heading estimation. Therefore, the set of surfaces is parsed due to inferred causal sources of motion and then channeled accordingly to the next processes.

In order to find a saddle point on a residual surface, we implemented saddle point operators designed to show activity when they are placed close to a saddle point area of a certain size and orientation. For that, the receptive field of such an operator consists of 5 circular same-sized areas arranged in a cross-shape. The operator averages the residual values in each area and compares the surrounding values to the central one. Operator activity is either the sum of absolute values of the differences between central and surrounding values or zero if the signs of those differences do not alternate. Hence, for an operator with non-zero activity there are exactly two directions from its central towards higher-valued areas, which we will call its peakward directions. In a multi-scale approach, we group saddle point operators that vary in the size of the circular areas of their receptive fields and their orientation and employ them throughout the incoming residual surface, which we transform by applying the negative logarithm and scaling the result to the range from 0 to 1. Summing the activity of each operator in such a group gives a cumulative activity for the retinal location at which that group was employed. This results in a retinotopically organized activity map that shows increased activity where the corresponding residual surface has a saddle point. An activity map with no or low activity indicates that the corresponding residual surface is due to observer flow only. Thus, if the activity map fails to exceed a certain *activity threshold τ*_1_, the residual surface will be assigned to the heading estimation layer or else to the object estimation layer.

The heading estimation layer sums all incoming surfaces, which results in the heading map. The peak in that map then indicates the estimated translational heading direction to explain the given flow field. As no further processing is needed beforehand, no operator type is employed for this layer.

The object estimation layer likewise aggregates all surfaces it receives. Then, to identify the location and properties of the IMO the saddle-point operators are applied again on the aggregated map. If enough activity is detected, that is, if the activity maximum surpasses the second activity threshold *τ*_2_, the model assumes the presence of an independent source of motion in the flow field. In that case, the retinal location of the maximum saddle-point activity is then declared as the estimated location of the object. The object direction estimate is a weighted average of peakward directions of the saddle point operators at the object’s estimated location. For every operator that contributes to the activity maximum, the direction with the smaller angle to the combined flow direction is used, weighted by the operator’s activity. By restricting the selection of the peakward directions to those more similar to the combined flow direction, we avoid opposing directions canceling out their contribution to the estimation. Additionally, the largest angle possible between the estimated direction and the object flow direction is 90°.

## 3. Model simulations

### 3.1. Paradigm

In the literature on self-motion and flow parsing a variety of paradigms have been employed to study various aspects of human perception, each well-designed to isolate a particular function or parameter set. Each of these paradigms would require a different set of settings and parameters for a model simulation (e.g. location, size, speed and direction of the IMO, speed of observer motion, size of the field of view, dot density, etc.). As we aim to show that our model is capable to simultaneously reproduce different facets of human behavior we decided to avoid implementing the individual specifics of each study and rather develop a generic simulation paradigm that allowed to compare the model behavior to essential findings across studies.

That paradigm establishes a self-motion scenario that consists of simulated observer translation towards a static cloud of dots together with an independently moving object. The cloud starts at 4m from the observer and has a depth of 6m, the object consists of a collection of dots in circular aperture at a particular location. These object dots are either all 4m from the observer or randomly placed inside the depth range the cloud covers. Dots in the cloud are placed so they are randomly distributed in the viewing window with height and width of 70 degrees of visual angle (dva) with 0.55dots/dva^2^. In this viewing window the object dots form a circular shape with a diameter of 1dva, 4dva or 8dva. Regardless of its size, the object always consisted of 50 dots. Background dots that are inside the objects shape are dismissed to make the object opaque. The object is placed relative to the translation direction of the observer and this offset is determined by its eccentricity to the FOE, ranging from 0dva to 15dva, in steps of 5dva, and one of 16 directions, starting from the one to the right in steps of 22.5° counter-clockwise.

Observer motion direction was randomly picked from the innermost 10dva of the FOV, the corresponding translation in the world *T* calculated, and the speed set to 2m/s. Object motion in the world consists of two separate components. The first is a horizontal translation *H* to the right of the observer, ranging from 0m/s to 1m/s, in steps of 0.125m/s. The second depends on the observer’s translation direction *T* and falls into one of three categories: (a) the object moves in the same direction as *T* so that it recedes from the observer, (b) the object actively approaches the observer by moving in the direction −*T* and (c) the second component is absent, making the object’s movement independent from the observer’s translation. The motion conditions are then defined by

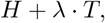

with *λ* ranging between 1 and 1, in steps of 0.5. This setup allows us to include the two motion conditions used typically in similar studies (Warren and Saunders, 1995; Royden and Hildreth, 1996; Layton and Fajen, 2016b; Li et al., 2018) to be the edge cases of our continuous set of motion conditions (s. Fig.3). We will refer to the motion conditions as ‘receding’, ‘semi-receding’, ‘neutral’, ‘semi-approaching’, and ‘approaching’, or with the corresponding values of *λ*. Similar to the study of Li et al. (2018), all object dots were placed at the front of the cloud for the ‘receding’ condition but varied in depth otherwise. Combinations of all the possible object offsets and sizes give rise to 147 different spatial layouts. We created five self-motion scenarios for each layout, then simulated observer motion and all object movements mentioned above for a total of 45 flow fields per scenario. Each flow field is then used as an input for the computational model. The heading space, the set of all candidate heading directions for which residual values are computed, covers the central 86dva × 86dva of the visual field. Candidates are sampled on a hexagonal grid so that adjacent directions are 1dva apart.

**Figure 3:**
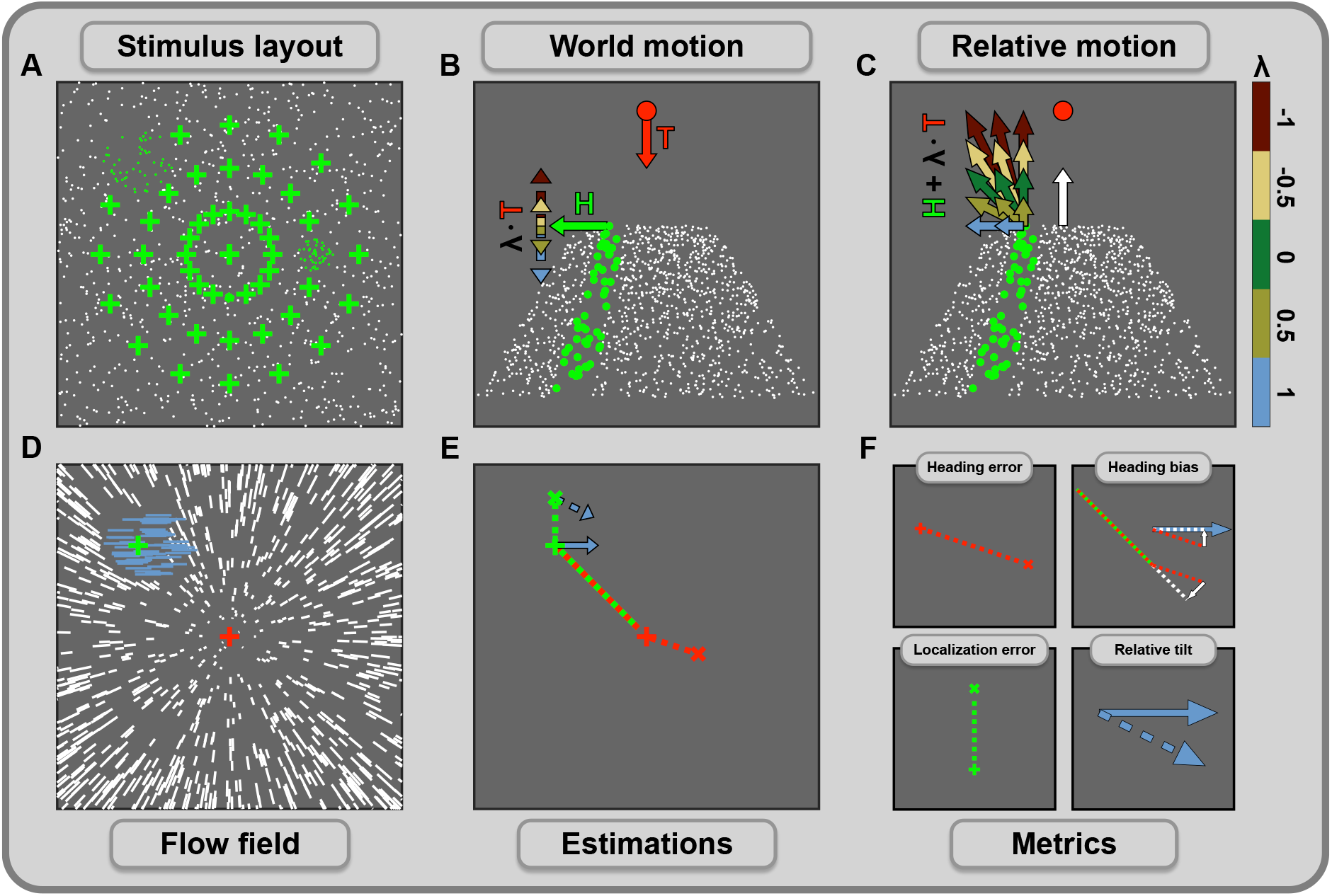
Simulation paradigm, estimation results, and performance metrics. (A) The object’s location in the visual field is determined relative to the direction of the observer’s movement. Potential object positions vary evenly in eccentricity and placement direction. While the simulation contains scenes with only one object present, panel A includes examples of three objects that vary in size and location. The green crosses indicate the remaining potential object positions, with the central one coinciding with the direction of observer movement. (B) The scene consists of the observer’s motion *T* toward a cloud of stationary dots. The self-moving object is opaque and occludes parts of the scene. Object movement consists of two components: a horizontal movement *H* (green arrow) at different speeds and a movement along the direction of the observer’s translation (*λ T*). This component defines the motion condition and ranges between the ‘receding’ (blue) and the ‘approaching’ (dark red) condition. (C) The motion in the scene is converted to relative motion between the world and the observer to compute the flow fields. (D) Flow fields are calculated for each combination of object size and location, the motion condition, and horizontal object speed. The example flow field shows the result of a simulated scene in the ‘receding’ condition. Combined flow is horizontal due to the backward motion canceling any changes in depth between the object and the observer. The red and green crosses indicate the observer’s translation direction and the object’s location, respectively. (E) The model estimates various scene parameters and whether the object is present for each flow field. The estimations of the heading direction (red x), the object location (green x), and the object’s movement direction (dashed blue arrow) are shown along the actual parameters. The green-red dashed line indicates the object offset relative to the heading direction. (F) Heading and localization error represent the distance between estimation and the true parameter in dva. Potential heading biases are indicated by the projection (solid white arrows) of the mis-estimation vector (red dashed line) onto the object direction and the offset vector. The estimated object direction is measured relative to the object flow direction by calculating the angle between them.

The model parameters were set to the following: First layer operators are placed on an evenly spaced rectangular grid so that the minimum distance to candidate directions is 0.5dva. Avoiding placing candidates and vectors of the flow representation too close reduces the possibility of irregular results during residual computation(Jepson and Heeger, 1990). Receptive fields of the first layer operators have a radius of 2dva. The receptive fields of the second layer operators have a radius of 20dva, are spaced 12dva apart and evenly placed to cover the FOV. This results in 2760 vectors in the optic flow field that is the output of the first layer and 36 residual surfaces calculated, one for each of the operators in the second layer. The groups of operators in the third layer that are placed on the residual surfaces are spaced 1dva apart. Sizes of the corresponding circular areas range from 1dva to 5dva in radius and the orientation of the cross-shape arrangement from 0° to 75°, relative to the cardinal axis. Activity threshold *τ*_1_ ranged between 0 and 6 and the detection threshold was set to *τ*_2_ = 1.5 *· τ*_1_.

### 3.2. Performance metrics

Certain metrics must be introduced to assess the model’s performance. A fundamental part of this model is the flow parsing that assigns residual surfaces to one of the subsequent layers. Measuring the quality of that assignment is crucial to determine the model’s capabilities because both processes, heading and object estimation, depend on the input to the corresponding layers. We evaluate *flow parsing quality* by calculating the rate at which our activity map criterion agrees with the actual causal source.

The following metrics examine the quality of the parameters that the model estimates. These metrics are visually depicted in Fig.2F. The primary metric to rate the heading estimation is the *heading error*, which is the distance between the true translation direction and the estimated translation direction in dva. Additionally, we calculate the extent to which the heading estimation error is in the direction of the object’s movement direction or its location, to unveil a potential *heading bias*. This is done by projecting the heading error vector onto the object direction and the offset vector, respectively.

Lastly, there are the metrics for the object estimation. We consider three aspects when measuring the object estimation performance of the model. For *object detection*, we check whether the binary estimation regarding the object’s presence correctly reflects the stimulus’s state. Similar to the heading estimation error, *localization error* is calculated as the distance between the estimated location and the retinal position of the object’s center. The estimated direction of the object’s movement is in retinal coordinates. We compare the actual direction of the combined flow with the estimated direction by calculating the angle between the corresponding vectors, resulting in the *relative tilt*. The sign of this angle reflects the direction in which the estimation is tilted, with positive values indicating a tilt in counter-clockwise direction and negative values a tilt in clockwise direction.

## 4. Results

### 4.1. Flow parsing

The main feature distinguishing our model from previous implementations of the subspace algorithm is that not all computed residual surfaces are used for heading estimation. Those residual surfaces that appear to be due to combined flow are used to extract information about the independent source of motion. The process for which a residual surface is used depends on its distribution and whether the maximum of the corresponding activity map surpasses the activity threshold *τ*_1_. Hence, the choice of *τ*_1_ regulates the flow parsing quality and, therefore, the input on which those processes depend. Thus, the first step in testing the model is to choose an appropriate threshold.

To determine a value for *τ*_1_, we started with a simulation in which the environment is entirely rigid. With no moving object in the scene, all flow is solely due to observer movement. The resulting residual surfaces are, therefore, all eligible for heading estimation. Fig.4A shows how the average rate of residuals used for heading estimation depends on the choice of the activity threshold. We want to adjust the flow parsing quality so that approximately 90% of surfaces enter the heading estimation layer. By design, raising the threshold increases the number of residuals that will be used for heading estimation. While it would be easily achievable to reach 100% by increasing the threshold value further, this would limit the input towards the object estimation layer when we include an IMO into the simulation. Hence, we set the activity threshold to *τ*_1_ = 3 for all following simulations as this puts the flow parsing at a level so that around 90% of residuals based solely on observer flow are used in the proper process.

**Figure 4:**
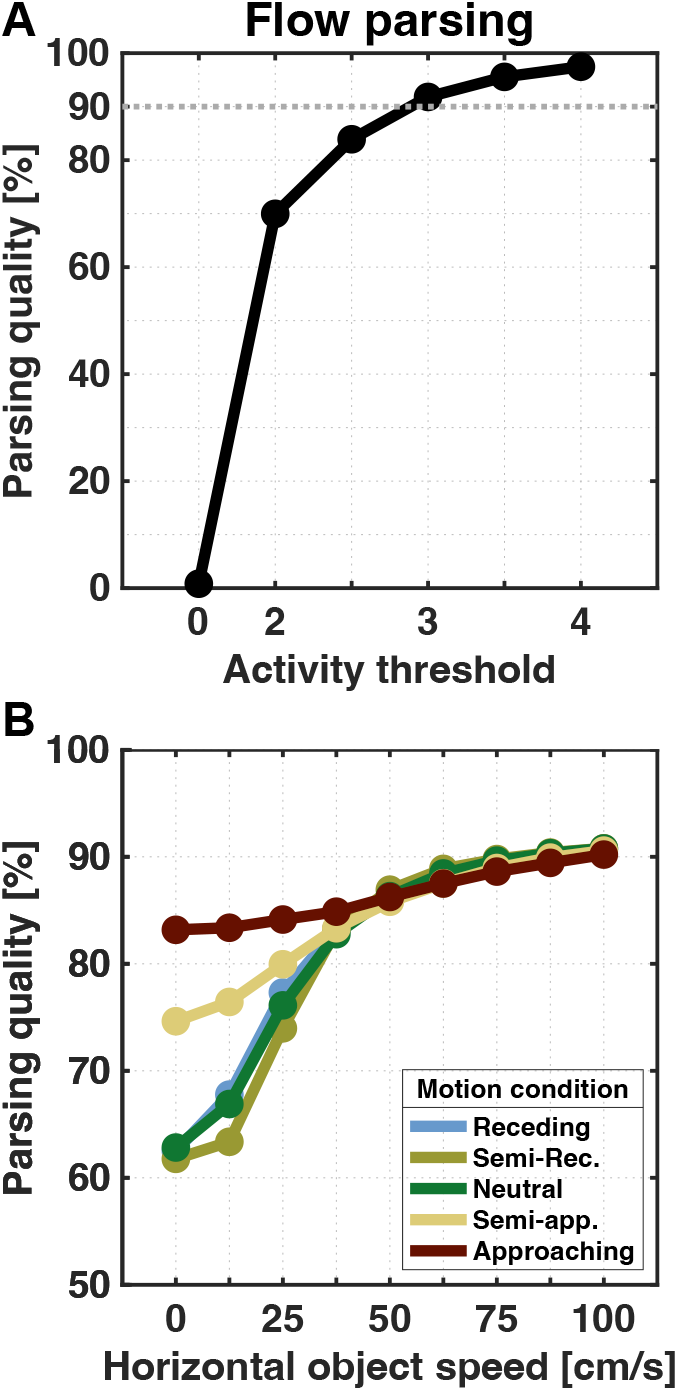
Flow parsing quality. Panel A shows the average rate of correctly assigned residual surfaces for different activity thresholds based on simulations without an IMO. Residuals whose activity maximum is below the threshold are used for heading estimation. The dashed line marks 90%, which we aim to surpass by our choice of the threshold. Panel B shows results for simulation with an IMO. Still, some residuals are only due to observer flow and should be assigned to the heading estimation layer. Lines indicate the average percentage rate at which the flow parsing process correctly parses the residuals into the respective layer.

For the main simulations we introduce an independent source of motion into the scene, so that some residuals are based on flow partly due to object movement and should, therefore, be used for object estimation. Fig.4B shows how often residuals end up in the correct layer and how this depends on the speed of the object’s horizontal movement and the motion condition. A maximum rate of around 90.8% is reached for all motion conditions when the object moves the fastest at 1*m/s*. On the other hand, the slowest horizontal object speed (0*m/s*) yields low rates of 61.8% − 62.8% for motion conditions in which the object is not approaching the observer. Otherwise, the residuals are correctly assigned at 74.6% and 83.2%, respectively. For all motion conditions, the increase in object speed lowers the chance of residuals being wrongly assigned for heading estimation.

For our simulation, we varied the horizontal speed of the object and found a strong influence on flow parsing quality. However, the object’s speed in the world is an ambiguous indicator for the combined flow’s velocity due to the various object locations and random depth distributions we tested. As the combined flow is the sum of observer flow and object flow, we can compute how adding the object’s movement changes the flow in speed and direction. While the speed ratio represents the quotient between the average speeds of the combined flow and the observer flow, the direction deviation is the average angle between the respective vectors of combined and observer flow. Computing those for each flow field allows us to present the relation between flow parsing quality and the speed ratio and direction deviation, as seen in Fig.6A. For flow fields that generated the lowest flow parsing quality, only 30.5% of residual surfaces were correctly assigned to the correct layer. In all these cases, the combined flow was either slower than the observer flow, so with a speed ratio below 1, or had a mean directional deviation below 10°. In general, the flow parsing quality increases when the combined flow is either faster or shows a higher deviation in direction. This is especially apparent when focussing on flow fields where combined flow only deviates in speed or direction (red dashed lines in Fig.6A). On average, 66.4% of residuals are correctly assigned for directional deviation below 10° and which rises to 83.8% when the average deviation is 45°. Similarly, average flow parsing quality is at 67.1% for speed ratios below 1 and at 82.8% when the speed ratio reaches 2. This shows, that the average flow parsing quality increases when the more the combined flow deviates from the pattern.

With the fixed threshold for flow parsing we can now turn to the other two tasks given to the model: heading estimation and object motion estimation. Our simulations are intended to show how the performance in these tasks depends on the parameters of the motion used in psychophysical studies.

### 4.2. Heading estimation

#### 4.2.1. Heading error

In order to estimate the direction of self-motion from an optic flow field, the model locates the peak of the heading map, which is the result of summing all residual maps assigned to the heading estimation layer. Fig.5A shows how close the estimation is to the actual parameter. When no self-moving object is in the scene, the heading error averages to 0.42dva. This is in line with human performance, as it is well established that humans can determine their heading from optic flow within 1 to 2 degrees of error when the flow is the result of simulated movement through a rigid scene (Warren and Hannon, 1988; Warren et al., 1988; Cutting et al., 1992). An independent source of motion in the scene disturbs the flow pattern due to observer movement. Thus, parts of the flow are no longer valid cues for self-motion. When an IMO is present in our simulation, the heading error follows a similar pattern regarding object speed for all motion conditions. When the object is not moving horizontally, the error is similar to baseline, ranging between 0.41dva and 0.62dva. It peaks at values between 0.88dva and 1.56dva for intermediate speeds before decreasing to 0.54dva for the fastest objects. The heading error is largest for motion conditions where the object recedes.

**Figure 5:**
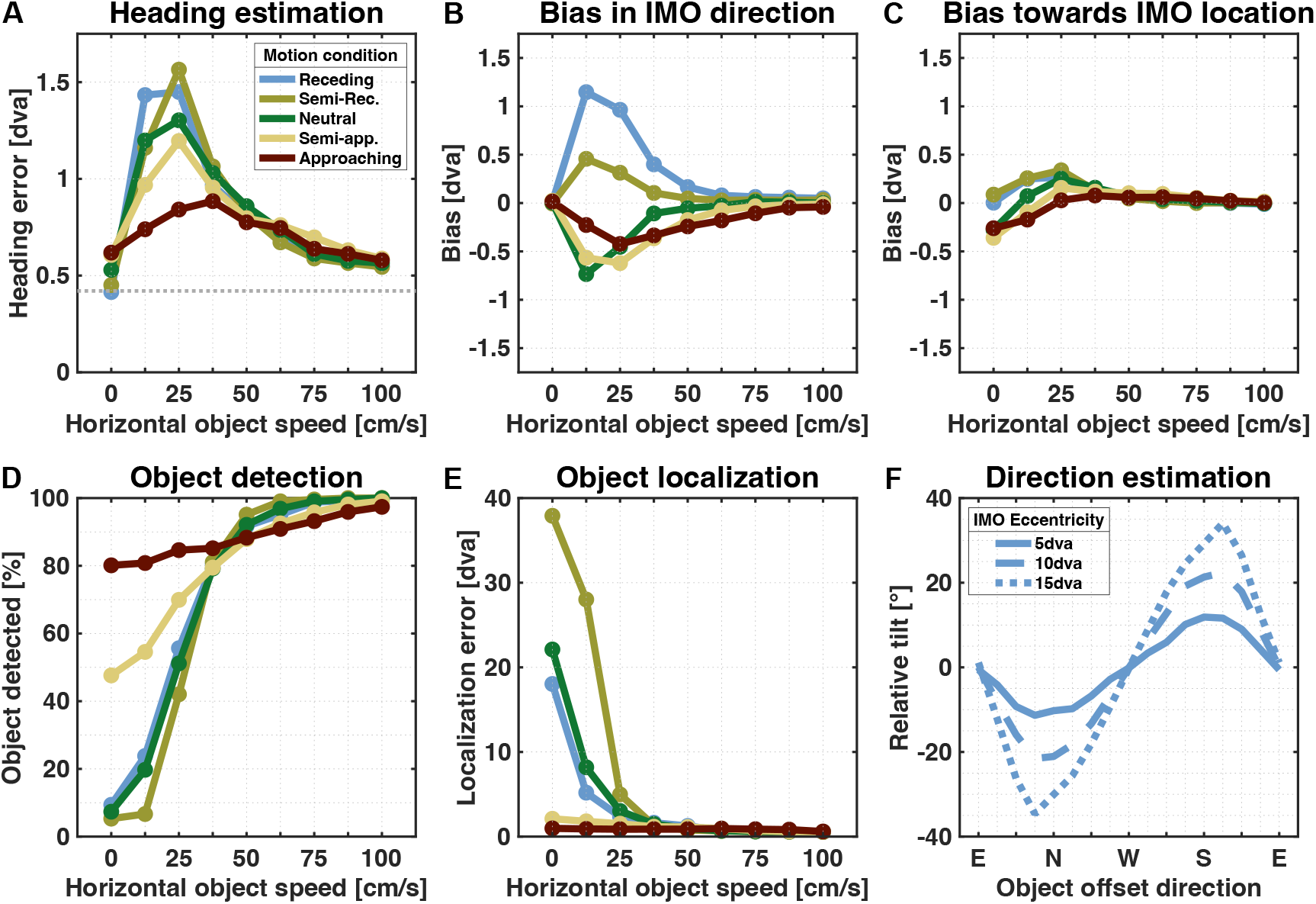
Heading and object estimation results. Panel A shows the heading error by horizontal object speed, averaged over all simulations. The dashed line represents the average error for flow fields without objects. Panels B and C show the heading error portion that is in the direction of object movement and position, respectively. Panel D represents the average object detection rate, and panel E contains the localization error for detected objects. Panel F shows the average angle between the estimated object movement direction and the combined flow direction for detected objects moving in the ‘receding’ condition. Results are presented for different object eccentricities, with positive values indicating an upward tilt and negative values indicating a downward tilt.

Interestingly, low flow parsing quality does not necessarily translate into an increased heading error. Residuals for a combined flow that is similar in direction to the observer flow are often wrongly attributed to the heading estimation pathway (Fig.6A). However, since the object direction is aligned with the direction of the observer flow these residuals are still a reasonably valid cue for heading estimation, such that only to a small error is produced in this situation (Fig.6B). On the other hand, for combined flow that deviates in direction from the observer flow, the heading error peaks at a direction change of 22° at 1.4dva. Deviation in direction from the observer flow increases the flow parsing quality and, thus, leads to a decrease in error. The largest heading error of 6.8dva was found for a flow field in which the combined flow was as fast as the observer flow at that location but deviated in direction by 48.5°.

**Figure 6:**
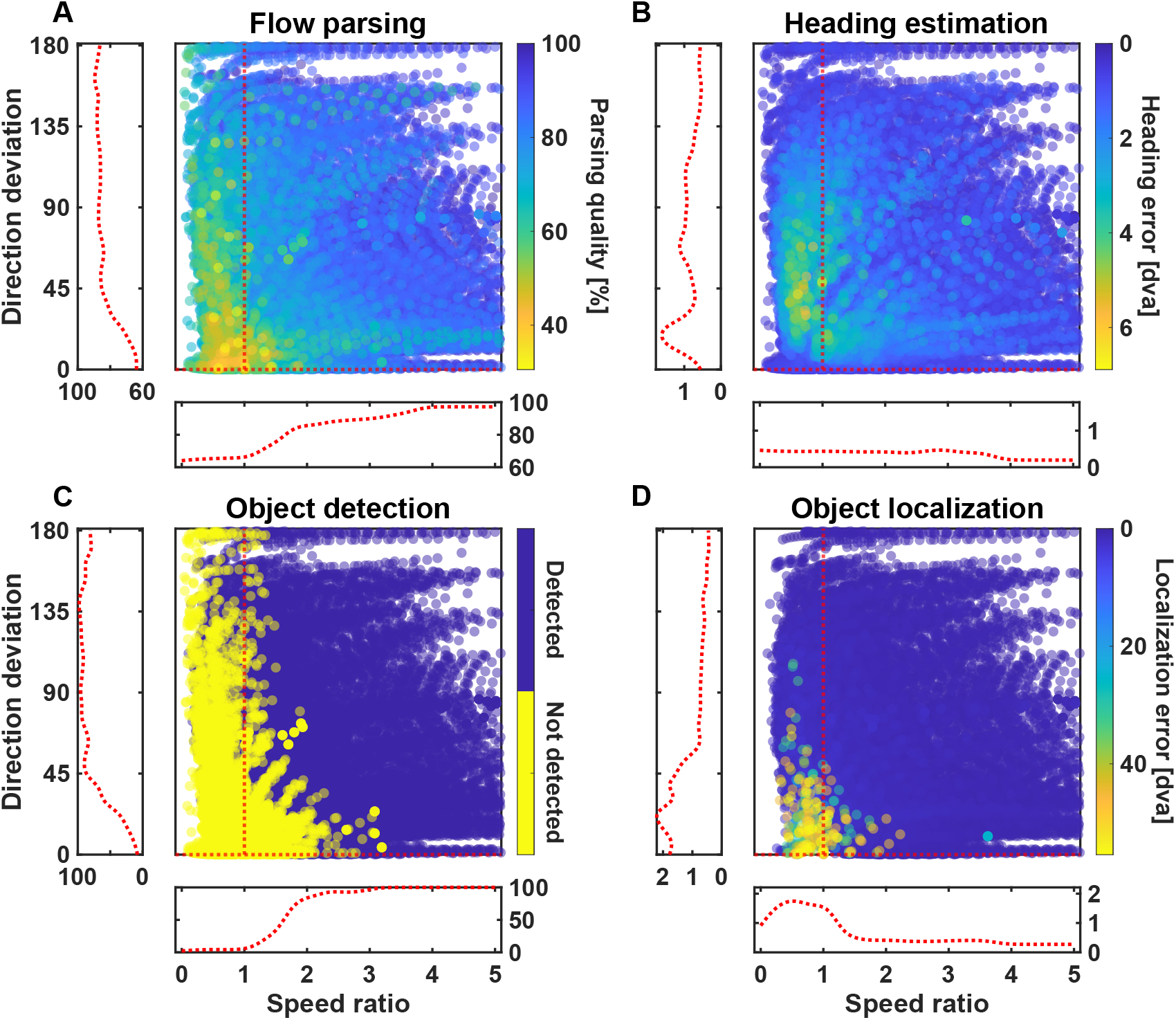
Performance by flow deviation. Results for all simulated flow fields are presented by speed ratio and directional deviation. Speed ratio is the quotient of the average speeds of the combined flow and observer flow at the object’s location. Values greater than 1 indicate that the combined flow is faster. Directional deviation is the average angle between the respective combined and observer flow vectors. The red dashed lines in the large panels indicate flow fields where the combined flow only deviates in either speed or direction. Running averages along those lines were computed to show the dependence of the performance on either of those aspects. To ensure enough data was available for such a computation, flow fields with combined flow deviating a maximum of 10% in speed and a maximum of 1° in direction, respectively, were included. Results can be seen to the left and below the large panels for the dependence of the performance on directional deviation and speed ratio, respectively.

The simulations of the present model revealed that a systematic variation of the object speed produced a particular pattern regarding the magnitude of the heading mis-estimation. Our results match the findings of the study of Dokka et al. (2019), who described that the dependence of heading error on object speed showed small mis-estimations for the slowest and fastest objects and peak error for intermediate speeds.

#### 4.2.2. Heading bias

To further characterize the systematic of the heading error, we calculated the extent of the misestimations in two particular directions: First, in the direction of the object’s movement, and second, in the direction towards its location. These potential biases can be seen in Fig.5B and Fig.5C. As the magnitude of a heading bias is limited by the size of the heading error, we focus on the range of object speeds yielding the largest mis-estimations, i.e., between 0.125m/s and 0.5m/s. When the object recedes, heading estimation is biased in the object’s direction for up to 1.15dva. For the other conditions, heading error is biased in the opposite direction of the object’s movement, peaking at 0.73dva. For the same range of speeds, we found a peak bias value of 0.34dva towards and 0.17dva away from the IMO location.

Comparing the peak bias values to the respective heading error, we can see that bias regarding object movement direction accounts for up to 80% of the heading mis-estimation. At the same time, bias towards or away from the IMO location covers a maximum of 23%.

We thus conclude, first, that a moving object can influence the perception of one’s own translation direction and, second, that the direction of that bias depends on the object’s movement direction and whether there is relative motion in depth between the object and the observer. These dependencies were found in several studies in human psychophysics (Warren and Saunders, 1995; Layton and Fajen, 2016c,b; Royden and Hildreth, 1996; Li et al., 2018). However, similar to our simulation, no such bias was seen towards the location of the object (Li et al., 2018).

### 4.3. Object estimation

#### 4.3.1. Object detection

Residual surfaces are assigned to the object estimation layer when they sport a saddle point pronounced enough that the maximum of the corresponding activity map exceeds *τ*_1_. The model assumes the presence of an IMO when the activity map we compute for the result of summing all of those incoming surfaces has a maximum that passes a second threshold *τ*_2_. By our choice of *τ*_1_, around 10% of residuals create enough activity to be misused for object estimation. Hence, based on the idea that the accumulation of residuals with saddle points pronounced enough to be assigned correctly for object estimation results in a surface with an even more distinct saddle point and, therefore, a higher activity maximum, we set *τ*_2_ = 1.5 *· τ*_1_.

With that, and as expected, the detection rate, as seen in Fig.5D, shows a similar pattern as the flow parsing quality (Fig.5A), namely that the faster objects are easier detected than slower ones. The detection rate rises above 97% for the fastest objects in all motion conditions. When there is no horizontal movement, the object is detected semi-reliably only when it approaches the observer, with detection rates of 47.6% and 80.1%, respectively. When the object is neither moving horizontally nor towards the observer, the detection rate is well below 10%. Only 5% of the objects moving such that the combined flow was as fast as or slower than the observer flow while in the same direction were detected. Objects that generated a higher speed ratio were detected more often, reaching detection rates of 50% and 75% at speed increases of 65% and 85%, respectively. These rates were also achieved by objects whose combined flow deviated by 27° and 41° but not in speed.

Studies showed that humans are indeed able to detect an IMO whose only cue was a disruption of a self-movement flow pattern, either in terms of a change in direction (Royden and Connors, 2010) or a change in flow speed (Royden and Moore, 2012). They found detection rates of 75% for directional deviations of around 15° and speed increases of around 40%.

#### 4.3.2. Object localization

While object detection is based on the value of the peak of the map computed in the object estimation layer, the object location is determined by the position of the peak. The localization error is the distance between the estimated location and the object’s center, if the IMO is detected. Hence, for objects with a radius of up to 4dva, estimated locations inside the object can still result in small localization errors. Results are shown in Fig.5E.

Approaching objects are localized reliably, regardless of horizontal speed. The average localization error was below 2.1dva in the ‘semi-approaching’ condition and below 0.97dva in the approaching’ condition. With no horizontal movement, the non-approaching objects were mislocalized with an average localization error between 18dva and 37.9dva. This error quickly drops with an increase in horizontal speed, going below 5dva when the objects move at 25cm/s and below 1dva at 62.5cm/s.

When the combined flow deviated only in speed, the average localization error dropped below 4dva at a speed increase of 85% due to the object movement. Similarly, a sole deviation in the flow direction of 46° led to an average localization error below 4dva.

The human object detection studies with which we compared our results in the previous section (Royden and Connors, 2010; Royden and Moore, 2012) only asked their participants to indicate if they noticed a self-moving object, but they were not asked to localize it. As we are unaware of other studies that included a localization task for IMOs in optic flow fields, we cannot compare our model’s performance to behavioral data. Thus, we can only conclude that one of our initial hypotheses, that the location of the emerging saddle point in a residual surface reliably indicates the object position, is valid since the localization of detected IMOs works remarkably well.

#### 4.3.3. Object direction estimation

The last aspect of the object estimation process is estimating the direction of the object’s movement. For this, we considered all objects that were detected and correctly localized in a radius of 10dva to the IMO center. To align with previous research in humans, we restrict our analysis to the ‘receding’ motion condition, where the combined flow was purely horizontal. Results can be seen in Fig.5F.

As the object in our paradigm is placed relative to the FOE, and this offset placement consists of the direction in which it is placed and its eccentricity, the findings can be easily summarized: Compared to the flow direction, the estimated direction is tilted downwards when the object is above the FOE but tilted upwards for objects below. The estimated direction is not tilted when the IMO is to the left or right of the FOE. Additionally, the closer the object is to the FOE, the smaller the effect. Peaks of the tilt are around 11.9°, 22.4°, and 34.6° in the corresponding direction for objects placed at 5dva, 10dva, and 15dva eccentricity, respectively.

Similar patterns were reported in studies where participants were asked to indicate an object’s trajectory in a flow field (Warren and Rushton, 2007, 2008, 2009; Rushton and Warren, 2005). The perceived movement direction was tilted towards the FOE compared to the on-screen motion, and the effect’s magnitude increased for objects farther away from the FOE. These responses were consistent with a scene-relative object motion. If the on-screen movement were interpreted as a result of self- and object motion, the tilt direction and its change in magnitude would match the subtraction of the flow component due to self-motion.

Due to the similar pattern in our estimations, we conclude that the direction our model estimates is the object’s world-relative movement direction in retinal coordinates rather than the direction of object flow.

### 4.4. Flow variations

Up to this point, the IMO in our simulations was surrounded by background points, and the flow we used as input for our model was either due to selfmotion or its combination with object movement. To test our model’s robustness, we deteriorated the flow fields by either introducing directional noise or presenting background flow in only one half of the visual field so that we could include the IMO in the other half. We focused on the’ receding’ condition to match existing psychophysical studies that used similar flow field variations. Additionally, we lowered the number of possible object offsets but kept the activity threshold *τ*_1_ the same as before.

#### 4.4.1. Directional noise

For this simulation, flow fields are computed as before. Before presenting them to our model, each background vector is altered by rotating it by a random degree, drawn from a Gaussian distribution, indicating the different noise levels we are testing. These are a ‘low noise’, a ‘mid noise’, and a ‘high noise’ condition characterized by standard deviations of 7.5°, 15° and 30°, respectively. The previous simulation is included as a ‘no noise’ condition.

In terms of flow parsing quality, adding noise lowers the rate at which residuals are correctly assigned (Fig.7). With no horizontal movement, the rate drops from 62.5% in the ‘no noise’ condition to around 38% for the three conditions that include noise. However, the improvement in flow parsing quality that comes with increased object speed is still present, with improvements of 21.7%, 14.4%, and 4.5%, depending on the noise level. Generally, the noise raises the overall activity computed from the respective residual surfaces. This leads to fewer residuals ending up in the heading estimation layer. Without noise, an average of 26.3% of the residuals are used for object estimation, which increases to 62.6%, 71.6%, and 88.1% for the three noise conditions, respectively.

**Figure 7:**
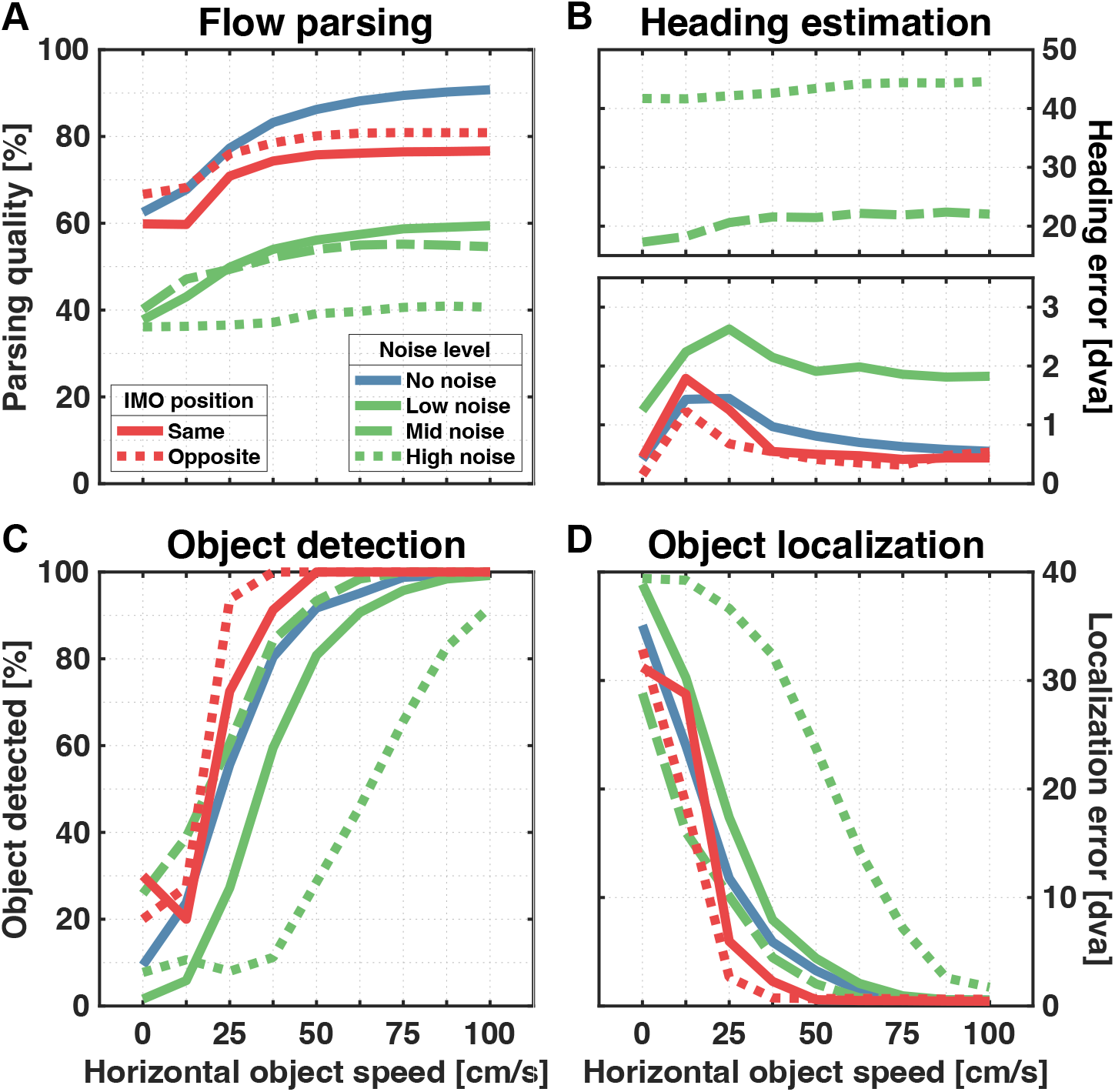
Estimation results for flow variations. Included are the results for the base simulation (blue lines, ‘receding’ condition), estimations for flow with added directional noise (green lines), and flow fields with background flow only in one hemifield relative to the object location (red lines). Panels show the flow parsing quality (A), the heading estimation error (B) and the performance in object detection (C) and localization (D).

Directional noise affects the heading estimation process in two ways. First, even pure self-motion flow has reduced reliability as a cue for heading. Second, as described above, fewer residual surfaces end up in the heading estimation layer. This leads to an increase in the average heading error to 1.1dva, 16.65dva, and 41.98dva for the respective noise conditions when no object is present and otherwise peaks at 2.63dva, 22.39dva, and 44.58dva. Furthermore, only the ‘low noise’ condition retains the previously found pattern (Fig.5B) in which the heading error increases at first before decreasing for faster object movement.

Contrary to the heading estimation layer, the object estimation layer receives a larger share of the residuals due to the noise, even though the number of residuals with object information does not change. Therefore, the object estimation results might be diluted due to wrongly assigned residuals. Regarding the overall pattern of the detection rate, it is kept intact as the rate rises with object speed, no matter the noise level (Fig.7C). Performance is lower in the ‘low noise’ and ‘high noise’ conditions as the object needs to be faster to reach the same detection rate as in the ‘no noise’ and ‘mid noise’ conditions. This indicates that the noise-induced increased activity for the individual residual surfaces does not directly translate to higher activity for the summed-up residual surface. Nonetheless, the ‘low’ and ‘mid’ conditions reach a detection rate of 99.2% and 100%, respectively. Even the ‘high noise’ condition surpasses 90% for the fastest objects.

The effect of noise on localization performance is similar to the one on detection rate. A steady improvement in the localization that comes with an increased object speed is present for all noise conditions (Fig.7D). While the ‘low noise’ and ‘mid noise’ conditions give rise to a localization error comparable to the ‘no noise’ condition, only the fastest objects are reliably localized when flow fields are confounded with the highest noise level.

The object direction estimation based on noisy flow shows the same tilt pattern for the simulation without noise (Fig.8A). It still holds that the estimated direction of objects placed above the FOE is tilted downwards, and vice versa, while for objects to the left or the right, there is no tilt on average. The magnitude of the peak tilt is reduced due to the noise, but only slightly for ‘low noise’ and ‘mid noise’ conditions with 19° and 21.6°, compared to the 22.7° in the ‘no noise’ condition. The highest noise level gives rise to the most prominent peak reduction down to 13.1°.

**Figure 8:**
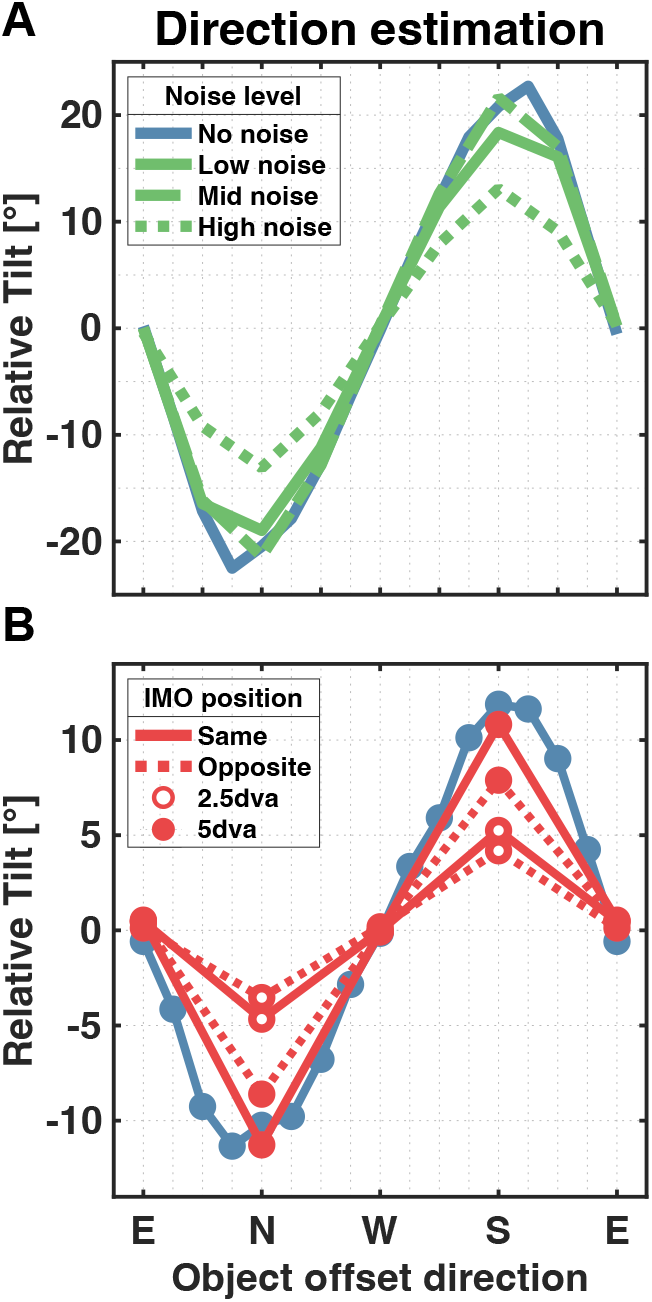
Object direction estimation for flow variations. Panels show the tilt of the estimated object direction compared to the horizontal combined flow based on the direction of the object offset. Panel A shows the tilt averaged over all object eccentricities and sizes tested for the different noise levels. Results were not averaged over object eccentricities in panel B as the base simulation did not include objects at an eccentricity of 2.5dva

These results are broadly consistent with several human studies that showed that self-motion estimation works reasonably well even when flow fields are confounded by various types of noise (Warren et al., 1991; Van Den Berg, 1992; Foulkes et al., 2013). The subspace algorithm, which forms a key part of our model, was also shown to be relatively robust when presented with noisy flow fields (Heeger and Jepson, 1992). Foulkes et al. (2013) showed that the algorithm is capable of producing similar patterns of estimated heading thresholds as humans, albeit with considerably higher average error. Regarding object direction estimation, Foulkes (2013) showed that, compared to noiseless flow fields, the introduction of noise decreased the peaks of the relative tilt pattern.

#### 4.4.2. Spatial isolation of the object

This simulation isolated the IMO from the remaining flow spatially, similar to a study by Warren and Rushton (2009). They aimed to test whether the assessment of world-relative motion from an independently moving object might be due to a local motion interaction between the object and the surrounding points. They removed the background flow in the hemifield the object was placed in and asked participants to indicate how they perceived the object’s trajectory. Results showed that the previously reported tilt pattern was still present but reduced.

For our simulation, observer translation was set to be towards the center of the FOV, and the object was positioned either above, below, or the left or the right of that. The object’s size was 1dva or 4dva in radius. Flow fields were computed as before but, background flow in one hemifield was removed, either so that the flow was on the ‘same’ side as the IMO or the ‘opposite’ side.

In terms of flow parsing quality, despite removing half of the background flow, the overall pattern is still intact as it increases alongside the object’s horizontal movement speed (Fig.7A). The rate of correct assignment of residuals plateaus at a lower level, around 76.5% and 80.9% for the ‘same’ and ‘opposite’ conditions, respectively, compared to the base simulation, which reaches up to 90.7%. However, this reduction might solely be due to a reduced number of residual surfaces assessed in the first place. Removing a hemifield of background flow led to fewer Layer 2 operators being necessary to cover the flow field, on average, a reduction of 15% based on the 36 operators used in the base simulation. Which, in turn, increased the rate of residuals partly due to combined flow by 7% up to 41%.

The heading estimation is quite similar to the base simulation, no matter which hemifield of flow was removed (Fig.7B). It peaks at 1.8dva and 1.2dva for objects moving at 12.5cm/s compared to the peak of 1.4dva at 25cm/s in the base simulation. For faster objects, the heading error drops to around 0.5dva in all three conditions.

Object detection and localization performance both show a slight improvement compared to the base simulation, as they already reach the limits of 100% detection rate and a localization error of around 0.5dva for objects moving 50cm/s compared to the 75cm/s needed in the base simulation (Fig.8C and D).

Similar to the previous results, presenting the model with only half the background flow does not break the tilt pattern previously found in the object direction estimations. The estimation remains tilted towards the FOE compared to the combined flow direction with a larger tilt when the IMO is placed at a higher eccentricity. Overall, the tilt was most prominent when the flow was presented in the whole FOV, peaking at 11.8°. Peak tilt was smaller at 11.3° and 5.2° when the object was placed in the ‘same’ hemifield as the flow at eccentricities of 5dva and 2.5dva, respectively. We found an even further decrease in tilt when the object was isolated from background flow with peak tilts of 8.6° and 4.2°.

Thus, the model results well match those reported by Warren and Rushton (2009).

## 5. Discussion

We presented a computational model to causally separate sources of motion in optic flow fields and recover information about self-motion and an independently moving object. The optic flow we used is represented by a vector flow field derived from a simulated self-motion scenario in which the object’s size, position, and velocity are systematically varied. Given an optic flow field as input, our model efficiently estimates the direction of self-motion, detects and locates an independent source of motion, and estimates the direction of its movement. Part of our model’s flow field evaluation process is to compute residual surfaces for flow from different parts of the visual field that indicate the consistency of the flow pattern with self-motion. Depending on the level of consistency, the surfaces are used either for heading or object estimation. While most of these aspects were already the subject of research over the years, studies often only covered one. To show that our model is capable of reproducing many of the reported behavioral patterns simultaneously, we developed a coherent simulation paradigm instead of recreating the stimuli used in all the different experiments.

### 5.1. Flow parsing and causal inference

The flow parsing hypothesis proposed that the brain uses its sensitivity to self-induced flow patterns to isolate object flow, following evidence that humans can judge the scene-relative motion of an object from retinal motion alone (Rushton and Warren, 2005; Warren and Rushton, 2007, 2008, 2009). While this process was not further specified, its description was often accompanied by the illustrative idea of “subtracting” a flow field due to self-motion from the retinal flow, because, despite the lack of plausibility for such an explicit subtraction process in the brain, it conceptually allows for easily understood predictions and reports. This has led to the thought that heading estimation is a prerequisite of flow parsing, since the flow component due to self-motion would need to be identified first, before it can be subtracted from the flow to reveal object motion. Rushton and colleagues have since clarified that this is not the case, and that flow parsing may not rely on prior heading estimation (Warren et al., 2012).

The flow parsing mechanism implemented in our model differs from the above approaches. But before pointing out the differences, we want to highlight the similarities. Most importantly, our implementation is a process that, among other things, assesses the scene-relative motion of an object while relying solely on visual input. In addition, it relies on sensitivity to flow patterns due to self-motion in the form of heading likelihood maps, which have already been used to model population responses of MST neurons (Lappe and Rauschecker, 1993; Lappe et al., 1996).

Our model differs from the original proposal in that we do not aim to isolate non-observer flow. Instead of working at the level of retinal motion, we propose a mid-level mechanism working on the population heading map. Our process infers the causal sources of the likelihood maps and, based on the result, passes them directly to the corresponding processes. The lack of a need to loop back to the retinal flow level for further processing may be a significant increase in processing speed. Furthermore, in our model heading estimation is not a prerequisite for flow parsing, but rather one of its results. This allows us to find the particular speed dependence of the heading error reported by Dokka et al. (2019). When the object moves slowly, the combined flow is mistakenly attributed to self-motion alone, even though it is not a valid cue. On the other hand, for faster objects, the flow is reliably recognized as being due to independent motion and used accordingly, reducing the impact on heading estimation performance.

Placing the flow parsing process at that mid-level stage of flow processing matches better to the results of studies in the recent years. Heading estimation is neither a prerequisite (Warren et al., 2012) nor limits the ability to assess scene-relative object motion (Rushton et al., 2018a). While directional noise directly affects the flow parsing quality by increasing the overall activity in layer 3, which impacts both types of processes, it additionally deteriorates the heading estimation performance. The reduced flow quality due to high levels of noise can render self-motion estimation ineffective but only lessens the tilt found in object direction estimation, leaving the overall pattern intact.

By separating the sources of retinal motion our flow parsing model can be seen as a process of causal inference. Traditionally, causal inference is modelled as a Bayesian integration process in which, for example, two sources of motion and their variances are optimally combined (e.g. Dokka et al. (2019)). While clearly many estimation processes in the visual system are Bayesian-optimal in this sense, such a view on causal inference does not address the question how heading and object motion are derived from the patterns of retinal motion. The Bayesian inference process initially requires knowledge of the true heading and object motion along with their measured variance. Our model differs from this approach by focusing on the computational mechanisms by which heading and object motion can be estimated directly from the retinal input. It is astounding that the results of this computation match the results of the Bayesian causal inference account (Dokka et al., 2019). It indicates that the computations of the model implement an optimal procedure for identifying separate sources of retinal motion.

### 5.2. Comparison to other computational models

Over the years, several heading estimation models have been proposed, some of which have taken on the challenge of explaining the results reported for scenes containing moving objects.

A prominent representative of motion pooling models is the model developed by Warren and Saunders (1995). Their two-layer model, which uses template matching to estimate the direction of self-motion, successfully replicated the heading estimation bias induced by approaching IMOs and explained it as the average of the FOEs due to observer and object motion. However, subsequent tests showed that it was not suitable for explaining the change in bias direction for purely lateral combined flow (Layton and Fajen, 2016a). Layton et al. (2012) followed a similar modeling approach, early motion pooling combined with template matching, but additionally equipped with competitive dynamics between the matching cells. While self-motion parameters were not estimated, heading biases were found in shifts of activity peaks of heading maps. These dynamics allowed to explain the change of heading bias direction for different types of object motion. Extending this model to include recurrent connections allowed it to capture the temporal dynamics of self-motion estimation in the presence of an IMO (Layton and Fajen, 2016a).

Another type of heading estimation model was inspired by the early work of Longuet-Higgins and Prazdny (1980) and Rieger and Lawton (1985). Their analysis of flow fields with rotational components showed that local differences in flow vectors, given a sufficient difference in depth, can provide a radial flow field for which the FOE coincides with the true heading direction. Hildreth (1992) used this method in a computational heading model so that it could deal with small, independently moving objects. By taking the direction that agrees with the majority of flow differences in different regions of the visual field as the estimated heading, small objects that caused inconsistent flow were omitted. Royden (1997) later adopted this idea and used motion-opponent operators inspired by neurons in primate area MT to implement this motion subtraction. Assuming a rigid scene apart from observer motion, heading was estimated by comparing maximally responding operators to translational heading templates. This model was later improved to deal with moving objects by adding Gaussian weighting to the connections between the operators of the first layer and the template cells (Royden, 2002). Interestingly, the author states that this addition was necessary to remove biases caused by objects away from the FOE, making this model less suitable for some of the newer data (Li et al., 2018). Another model extension resulted in one of the few optic flow-based models for object detection (Royden and Holloway, 2014). After heading estimation, the first layer operators’ preferred directions and response magnitudes were compared to the template that determined the estimate. If the direction differed too much or the response magnitude was significantly higher than the responses of other cells, it was assumed that there was a self-moving object in the scene with a boundary at the operator’s location. While this model showed promising and robust results under various circumstances, it used heading estimation as a prerequisite for the object detection process.

The approach Raudies and Neumann (2013) used in their computational study to estimate selfmotion from optic flow containing an IMO was different. Their analytic model relied on local segmentation cues such as accretion/deletion, expansion/contraction, and acceleration/deceleration to qualitatively reproduce the behavioral pattern of heading bias. This, however, is not in line with the results of Li et al. (2018) who showed that the heading estimation process does not include the segmentation of independently moving objects.

Compared to optic flow based heading estimation models, the field of object motion estimation models is sparsely populated. Layton and Fajen (2016b) presented a neurophysiologically inspired model of object motion recovery during self-motion. It uses interactions in MT and feedback from MSTd to MT to transform retinal object motion into a world-relative reference frame. A key prediction was that such a process, which shifts initial MT responses reflecting the retinal motion pattern to align with world-relative motion, depends on a temporal process in MT. A more elaborate version of the model has been developed by Layton and Niehorster (2019). It uses two separate processing streams, one for self-motion estimation consisting of MT cells with reinforcing surrounds projecting to MSTd, and the other using MT cells with suppressive surrounds connected to ventral MST, to estimate scene-relative object motion. The latter pathway is modulated by the estimates of the former. While recent findings indicate that MT activity is modulated in accordance with scene-relative object motion (Peltier et al., 2024), it remains open whether the reported time course matches the time course of real life situations, in which eye movements frequently disrupt the retinal flow such that only segments of a duration of about 300ms are available for processing (Matthis et al., 2022).

The subspace algorithm, published by Heeger and Jepson (1992) and implemented in a heading estimation model, recovers self-motion parameters under the assumption of a rigid scene. Potential biases due to object motion would result from systematic mis-estimation of self-motion. Lappe and Rauschecker (1993), who designed their population heading map model by using this algorithm to determine connection weights between neural layers, found that the most consistent interpretation of a flow field containing purely lateral combined flow was a combination of observer translation and rotation. The estimated direction was shifted in the direction of object motion to counteract rotation for the rest of the flow field (Li et al., 2018). On the other hand, the explanation for an approaching object was a translation with an offset angle. Hence, the model shifted the estimate in the opposite direction of the object’s motion. Since our model also uses the subspace algorithm for heading estimation, the biases we found can also be explained in this way. While our implementations of the various flow processing steps may be lacking in the neurophysiological formulation, we want to emphasize that Lappe and Rauschecker (1993) have already covered the first stages of our model by computing a plausible representation of the incoming flow and the residual surfaces. Furthermore, the process of flow parsing and subsequent estimation consists, in simple terms, of counting and localizing peaks on these retinotopic surfaces. This would remain the last hurdle in transforming our model into a neurophysiologically plausible one.

### 5.3. Summary

This detailed discussion of experimental results and different types of models aimed to show the unique position in which our model fits. While it provides different types of estimations, ranging from self-motion to object estimation, it reproduces behavioral trends found in various studies. The key to that is the flow parsing process we implemented, which is based on sensitivity to optic flow patterns consistent with self-motion. Our interpretation of the flow parsing process works on the residual surfaces computed for parts of the optic flow and does not parse the flow itself into its components. While this could be achieved retrospectively, it is not necessary in the context of our model, as estimations of all the parameters are based on these surfaces and not isolated flow. Additionally, the straightforward structure of the model shows that recurrent connections or feedback loops might not be necessary for those processes. In our model, heading estimation and independent object motion estimation work in parallel and in a fast, feedforward fashion.

## Notes

### Competing Interest Statement

The authors have declared no competing interest.

### Summary of Updates

Corrected Pabel 4 of Figure 6. Reason: Used wrong data (wrong: Localization of all objects, intended: Localization of all detected objects) Clarified in the text that localization error refers to the localization performance of detected objects.

## References

Albright, T. D. (1984). Direction and orientation selectivity of neurons in visual area MT of the macaque. Journal of Neurophysiology, 52(6):1106–1130.

Calow, D., Krüger, N., Wörgotter, F., and Lappe, M. (2005). Biologically motivated space-variant filtering for robust optic flow processing. Network: Computation in Neural Systems, 16(4):323–340.

Calow, D. and Lappe, M. (2007). Local statistics of retinal optic flow for self-motion through natural sceneries. Network, 18(4):343–374.

Cutting, J. E., Springer, K., Braren, P. A., and Johnson, S. H. (1992). Wayfinding on foot from information in retinal, not optical, flow. Journal of Experimental Psychology: General, 121(1):41–72.

Dokka, K., Park, H., Jansen, M., DeAngelis, G. C., and Angelaki, D. E. (2019). Causal inference accounts for heading perception in the presence of object motion. Proceedings of the National Academy of Sciences, 116(18):9060–9065.

Duffy, C. J. and Wurtz, R. H. (1991). Sensitivity of MST neurons to optic flow stimuli. I. A continuum of response selectivity to large-field stimuli. Journal of Neurophysiology, 65(6):1329–1345.

Foulkes, A. (2013). Flow parsing and heading perception show similar dependence on quality and quantity of optic flow. Frontiers in Behavioral Neuroscience, 7.

Foulkes, A. J., Rushton, S. K., and Warren, P. A. (2013). Heading recovery from optic flow: comparing performance of humans and computational models. Frontiers in Behavioral Neuroscience, 7.

Gibson, J. J. (1950). The perception of the visual world. Houghton Mifflin, Boston.

Gray, R., Macuga, K., and Regan, D. (2004). Long range interactions between object-motion and self-motion in the perception of movement in depth. Vision Research, 44(2):179–195.

Gray, R. and Regan, D. (2000). Simulated self-motion alters perceived time to collision. Current Biology, 10(10):587– 590.

Grigo, A. and Lappe, M. (1999). Dynamical use of different sources of information in heading judgments from retinal flow. Journal of the Optical Society of America A, 16(9):2079.

Heeger, D. J. and Jepson, A. D. (1992). Subspace methods for recovering rigid motion I: Algorithm and implementation. International Journal of Computer Vision, 7(2):95– 117.

Hildreth, E. C. (1992). Recovering heading for visually-guided navigation. Vision Research, 32(6):1177–1192.

Jepson, A. D. and Heeger, D. J. (1990). Subspace methods for recovering rigid motion II: Theory. Technical Report RBCV-TR-90-36, University of Toronto.

Lappe, M. (1996). Functional Consequences of an Integration of Motion and Stereopsis in Area MT of Monkey Extrastriate Visual Cortex. Neural Computation, 8(7):1449– 1461.

Lappe, M. (1998). A model of the combination of optic flow and extraretinal eye movement signals in primate extrastriate visual cortex Neural model of self-motion from optic flow and extraretinal cues. Neural Networks.

Lappe, M., Bremmer, F., Pekel, M., Thiele, A., and Hoffmann, K. P. (1996). Optic flow processing in monkey STS: a theoretical and experimental approach. J. Neurosci., 16(19):6265–6285.

Lappe, M., Bremmer, F., and van den Berg, A. (1999). Perception of self-motion from visual flow. Trends in Cognitive Sciences, 3(9):329–336.

Lappe, M., Pekel, M., and Hoffmann, K. P. (1998). Optokinetic eye movements elicited by radial optic flow in the macaque monkey. J. Neurophysiol., 79(3):1461–1480.

Lappe, M. and Rauschecker, J. P. (1993). A Neural Network for the Processing of Optic Flow from Ego-Motion in Man and Higher Mammals. Neural Computation, 5(3):374– 391.

Lappe, M. and Rauschecker, J. P. (1994). Heading detection from optic flow. Nature, 369(6483):712–713.

Lappe, M. and Rauschecker, J. P. (1995a). An illusory transformation in a model of optic flow processing. Vision Research, 35(11):1619–1631.

Lappe, M. and Rauschecker, J. P. (1995b). Motion anisotropies and heading detection. Biological Cybernetics, 72(3):261–277.

Layton, O. W. and Fajen, B. R. (2016a). Competitive Dynamics in MSTd: A Mechanism for Robust Heading Perception Based on Optic Flow. PLOS Computational Biology, 12(6):e1004942.

Layton, O. W. and Fajen, B. R. (2016b). A Neural Model of MST and MT Explains Perceived Object Motion during Self-Motion. Journal of Neuroscience, 36(31):8093–8102.

Layton, O. W. and Fajen, B. R. (2016c). Sources of bias in the perception of heading in the presence of moving objects: Object-based and border-based discrepancies. Journal of Vision, 16(1):9.

Layton, O. W. and Fajen, B. R. (2016d). The temporal dynamics of heading perception in the presence of moving objects. Journal of Neurophysiology, 115(1):286–300.

Layton, O. W., Mingolla, E., and Browning, N. A. (2012). A motion pooling model of visually guided navigation explains human behavior in the presence of independently moving objects. Journal of Vision, 12(1):20–20.

Layton, O. W. and Niehorster, D. C. (2019). A model of how depth facilitates scene-relative object motion perception. PLOS Computational Biology, 15(11):e1007397.

Li, L., Ni, L., Lappe, M., Niehorster, D. C., and Sun, Q. (2018). No special treatment of independent object motion for heading perception. Journal of Vision, 18(4):19.

Li, L. and Warren, W. H. (2000). Perception of heading during rotation: sufficiency of dense motion parallax and reference objects. Vision Research, 40(28):3873–3894.

Longuet-Higgins, H. C. and Prazdny, K. (1980). The interpretation of a moving retinal image. Proceedings of the Royal Society of London. Series B, Biological Sciences, 208(1173):385–397.

Matthis, J. S., Muller, K. S., Bonnen, K. L., and Hayhoe, M. M. (2022). Retinal optic flow during natural locomotion. PLoS Comput Biol, 18(2):e1009575.

Maunsell, J. H. and Van Essen, D. C. (1983). Functional properties of neurons in middle temporal visual area of the macaque monkey. I. Selectivity for stimulus direction, speed, and orientation. Journal of Neurophysiology, 49(5):1127–1147.

Pauwels, K., Krüger, N., Lappe, M., Wörgötter, F., and Van Hulle, M. M. (2010). A cortical architecture on parallel hardware for motion processing in real time. Journal of vision, 10(10):18–18.

Peltier, N. E., Anzai, A., Moreno-Bote, R., and DeAngelis, G. C. (2024). A neural mechanism for optic flow parsing in macaque visual cortex. Current Biology, page S0960982224012417.

Raudies, F. and Neumann, H. (2013). Modeling heading and path perception from optic flow in the case of independently moving objects. Frontiers in Behavioral Neuroscience, 7.

Regan, D. and Beverley, K. I. (1982a). How do we avoid confounding the direction we are looking and the direction we are moving? Science, 215:194–196.

Regan, D. and Beverley, K. I. (1982b). How Do We Avoid Confounding the Direction We Are Looking and the Direction We Are Moving? Science, 215(4529):194–196.

Rieger, J. H. and Lawton, D. T. (1985). Processing differential image motion. Journal of the Optical Society of America A, 2(2):354.

Royden, C. S. (1997). Mathematical analysis of motion-opponent mechanisms used in the determination of heading and depth. Journal of the Optical Society of America A, 14(9):2128.

Royden, C. S. (2002). Computing heading in the presence of moving objects: a model that uses motion-opponent operators. Vision Research, 42(28):3043–3058.

Royden, C. S., Banks, M. S., and Crowell, J. A. (1992). The perception of heading during eye movements. Nature, 360(6404):583–585.

Royden, C. S. and Connors, E. M. (2010). The detection of moving objects by moving observers. Vision Research, 50(11):1014–1024.

Royden, C. S., Crowell, J. A., and Banks, M. S. (1994). Estimating heading during eye movements. Vision Research, 34(23):3197–3214.

Royden, C. S. and Hildreth, E. C. (1996). Human heading judgments in the presence of moving objects. Perception & Psychophysics, 58(6):836–856.

Royden, C. S. and Holloway, M. A. (2014). Detecting moving objects in an optic flow field using direction- and speed-tuned operators. Vision Research, 98:14–25.

Royden, C. S. and Moore, K. D. (2012). Use of speed cues in the detection of moving objects by moving observers. Vision Research, 59:17–24.

Rushton, S. K., Bradshaw, M. F., and Warren, P. A. (2007). The pop out of scene-relative object movement against retinal motion due to self-movement. Cognition, 105(1):237–245.

Rushton, S. K., Chen, R., and Li, L. (2018a). Ability to identify scene-relative object movement is not limited by, or yoked to, ability to perceive heading. Journal of Vision, 18(6):11.

Rushton, S. K., Niehorster, D. C., Warren, P. A., and Li, L. (2018b). The primary role of flow processing in the identification of Scene-Relative object movement. J Neurosci, 38(7):1737–1743.

Rushton, S. K. and Warren, P. A. (2005). Moving observers, relative retinal motion and the detection of object movement. Current Biology, 15(14):R542–R543.

Sauer, Y., Scherff, M., Lappe, M., Rifai, K., Stein, N., and Wahl, S. (2022). Self-motion illusions from distorted optic flow in multifocal glasses. iScience, 25(1):103567.

Van Den Berg, A. (1992). Robustness of perception of heading from optic flow. Vision Research, 32(7):1285–1296.

Warren, P. A. and Rushton, S. K. (2007). Perception of object trajectory: Parsing retinal motion into self and object movement components. Journal of Vision, 7(11):2.

Warren, P. A. and Rushton, S. K. (2008). Evidence for flow-parsing in radial flow displays. Vision Research, 48(5):655–663.

Warren, P. A. and Rushton, S. K. (2009). Perception of scene-relative object movement: Optic flow parsing and the contribution of monocular depth cues. Vision Research, 49(11):1406–1419.

Warren, P. A., Rushton, S. K., and Foulkes, A. J. (2012). Does optic flow parsing depend on prior estimation of heading? Journal of Vision, 12(11):8–8.

Warren, W. H., Blackwell, A. W., Kurtz, K. J., Hatsopoulos, N. G., and Kalish, M. L. (1991). On the sufficiency of the velocity field for perception of heading. Biological Cybernetics, 65(5):311–320.

Warren, W. H. and Hannon, D. J. (1988). Direction of self-motion is perceived from optical flow. Nature, 336(6195):162–163.

Warren, W. H. and Hannon, D. J. (1990). Eye movements and optical flow. Journal of the Optical Society of America A, 7(1):160.

Warren, W. H., Morris, M. W., and Kalish, M. (1988). Perception of translational heading from optical flow. Journal of Experimental Psychology: Human Perception and Performance, 14(4):646–660.

Warren, W. H. and Saunders, J. A. (1995). Perceiving Heading in the Presence of Moving Objects. Perception, 24(3):315–331.

